# Markers of mitochondrial function and oxidative metabolism in female skeletal muscle do not display intrinsic circadian regulation

**DOI:** 10.1101/2025.01.08.631231

**Authors:** Liam S. Fitzgerald, Connor S. Reynoso Spurrier, Nathan J. Lau, Miles D. Melamed, Lindsey A. Burnett, Gretchen A. Meyer, Chang Gui, Andrea L. Hevener, James A. Sanford, Simon Schenk

## Abstract

Mitochondria are key regulators of metabolism and ATP supply in skeletal muscle, while circadian rhythms influence many physiological processes. However, whether mitochondrial function is intrinsically regulated in a circadian manner in mouse skeletal muscle is inadequately understood. Accordingly, we measured post-absorptive transcript abundance of markers of mitochondrial biogenesis, dynamics, and metabolism (extensor digitorum longus [EDL], soleus, gastrocnemius), protein abundance of electron transport chain complexes (EDL and soleus), enzymatic activity of SDH (tibialis anterior and plantaris), and maximum uncoupled respiration (tibialis anterior) in different skeletal muscles from female C57BL/6NJ mice at four zeitgeber times (ZT), ZT 1, 7, 13, and 19. Our findings demonstrate that markers of mitochondrial function and oxidative metabolism do not display intrinsic time-of-day regulation at the gene, protein, enzymatic, or functional level. The core-clock genes *Bmal1* and *Dbp* exhibited intrinsic circadian rhythmicity in skeletal muscle (i.e., EDL, soleus, gastrocnemius) and circadian amplitude varied by muscle type. These findings demonstrate that female mouse skeletal muscle does not display circadian regulation of markers of mitochondrial function or oxidative metabolism over 24 hours.

## Introduction

Circadian rhythms are self-sustaining oscillations in an organism or cell that repeat approximately every 24 hours. In mammalian physiology, for example, blood pressure (1), body temperature (2), and hormone secretion (3), are well-known to exhibit circadian rhythmicity. More recently, whether other aspects of mammalian physiology and metabolism exhibit circadian oscillation has been intensely investigated (4–13). To this end, because mitochondria are fundamental regulators of physiology, metabolism, and function in skeletal muscle, there is great interest in understanding the impact of time-of-day on mitochondrial morphology, homeostasis, and function (13–17). For example, time-of-day variation in muscle performance during exercise is proposed to be regulated by circadian (i.e., time-of-day) changes in mitochondrial function (i.e., oxidative capacity) (13, 18). Thus, the prevailing belief in the field is that mitochondrial function in skeletal muscle is intrinsically regulated (i.e., ‘programmed’) to oscillate by time-of-day (19–21).

Six lines of evidence have helped establish the perspective that mitochondrial function in skeletal muscle is intrinsically regulated by time-of-day; specifically, studies in 1) immortalized C2C12 cells (22), 2) primary mouse myoblast cell lines (23), 3) mice with germline global knockout of core-clock genes (24–26), 4) mice with muscle-specific knockout of core-clock genes (27), 5) rats (28, 29), and 6) humans (20, 30, 31). For example, in C2C12 myoblasts, oxygen consumption oscillates over a 24-h period, independently of serum exposure, suggesting intrinsic 24-h cycles of mitochondrial oxidative metabolism (22). In mice with whole-body knockout of the core-clock gene *Bmal1* (Bmal1^-/-^), decreased mitochondrial volume and a lower respiratory control ratio (state III/state IV) was observed in gastrocnemius and diaphragm compared to wildtype (WT) (24). Moreover, maximal fatty acid oxidation is higher in primary myotubes established from *Cry1*^*−/−*^*;Cry2*^*−/−*^ double knockout mice (dKO^*Cy1/Cry2*^) compared to WT littermates (23). The effect of time-of-day on mitochondrial function, however, is not a universal finding as no effect of time-of-day on maximal mitochondrial respiration was found in skeletal muscle from humans (20, 32) or *ad libitum*-fed rats (29).

Overall, the generalized thought in the field is that markers of mitochondrial function are intrinsically regulated by time-of-day in skeletal muscle. However, if markers of mitochondrial function are intrinsically regulated by time-of-day, then by definition time-of-day-dependent oscillations in markers of mitochondria function should be shared (although perhaps of different magnitude) across different skeletal muscles. To our knowledge, however, the impact of time-of-day on markers of mitochondrial function has not been rigorously studied in multiple muscles or when controlling for timing of food intake, which profoundly impacts skeletal muscle gene expression (34–36) and metabolism (51, 29). Further, circadian rhythm studies on mitochondrial function in skeletal muscle have only studied male rodents (5, 12, 22, 23, 25, 26, 32, 38, 48, 29, 33) and almost exclusively male humans (20, 30, 50, 31). Addressing these gaps, in this study we measured markers of mitochondrial function at the gene, protein, or enzymatic level in several different skeletal muscles from female mice. Specifically, we assessed extensor digitorum longus (EDL), soleus (SOL), gastrocnemius (GA), tibialis anterior (TA), plantaris (PLN), at 4 different times-of-day (namely, zeitgeber time [ZT] ZT 1, 7, 13, and 19). Further, all mice were given a standardized last meal 3 hours before tissue collection.

Based on our previous work demonstrating that SOL and EDL endurance capacity, as a functional marker of mitochondrial function, is not intrinsically regulated by time-of-day in female or male mice (37), and the fact that mitochondrial terms were not enriched in the seminal skeletal muscle circadian transcriptome papers by McCarthy et al. (26) and Miller et al. (38), we hypothesized that markers of mitochondrial function and oxidative metabolism in female mouse skeletal muscle would not exhibit intrinsic circadian regulation.

## Materials and Methods

### Animals

The animals used in this study have been described previously (37). Briefly, 13.0±0.1-week-old female C57BL/6NJ mice (The Jackson Laboratory, stock #05304) were studied. For 3 weeks before the studies, animals were co-housed (3-5 mice/cage) on a 12-hour light/12-hour dark cycle – Lights on: 06:00 h (ZT 0); Lights off (Dark): 18:00 h (ZT 12). There were four experimental groups: ZT 1, 7, 13, 19, with ZT1 and ZT 13, and ZT 7 and ZT 19, chosen to represent “early” and “late” timepoints within each light/dark cycle, respectively. Experiments during the dark phase were performed under dim red light with tissue dissection occurring after anesthetization, under normal ambient light. To account for potentially confounding effects of nutritional status on gene expression and mitochondrial function across time-of-day (33), mice were gavaged with a standardized dextrose ‘meal’ (2 g/kg), 3 hours before tissue dissection; they were then fasted with *ad libitum* access to water. All animal experiments were approved by and conducted in accordance with the Animal Care Program at the University of California, San Diego.

### Tissue Dissection

After an intraperitoneal injection of a pentobarbital/phenytoin-containing solution (300 mg/kg; Euthasol; Virbac), the EDL, SOL, GA, PLN, and TA were rapidly dissected. EDL, SOL, and GA were blotted dry and immediately frozen in liquid nitrogen while PLN and TA were pinned at a physiologic muscle length and frozen in liquid nitrogen-cooled isopentane. Samples were stored at −80°C for subsequent analysis.

### RNA extraction, reverse transcription, and real-time PCR

RNA was extracted with TRIzol (Invitrogen) and RNA concentration and quality was measured (NanoDrop 2000 spectrophotometer), with 500ng of RNA used for cDNA synthesis (Applied Biosystems). Semi-quantitative real-time PCR analysis was conducted on 50 ng of cDNA using SYBR Green master mix (Thermo Scientific).

Relative expression for each gene was calculated using the ΔΔCt method, with Ribosomal Protein Lateral Stalk Subunit P0 (*Rplp0*) expression as the normalization control. For each gene, the time-point with the lowest expression level (the nadir) was averaged to 1.0 and all other time-points were expressed relative to that point. The primers are listed in **Supplementary Table 1**.

### Muscle homogenization, capillary-based gel electrophoresis and immunodetection

Muscles were homogenized and the lysate isolated, as previously described (37). The Jess System (Protein Simple) was used for immunodetection (4 μg protein), as previously described (37). The primary antibodies, OxPhos complex kit (Invitrogen, # 458099; 1/75 dilution) and eEF2 (Cell Signaling, #2332; 1/50 dilution) were run together in each capillary. Secondary probing was conducted with a 50:50 mix of anti-rabbit (Bio-Techne, #042-206) and anti-mouse secondary HRP antibodies (Bio-Techne, #042-205). Proteins were normalized to eEF2.

### Succinate dehydrogenase (SDH) activity

Pinned plantaris and tibialis anterior were embedded in OCT (Fisher), sectioned (10 μm) and mounted onto 25×75 microscope slides. Slides were incubated in reaction solution (1.5 mM nitroblue tetrazolium, 6.4 mM EDTA, 110 mM succinic acid, 0.75 mM sodium azide, 30 mM 1-methoxyphenazine methosulphate, 13 mM KH_2_PO_4_, and 85 mM K_2_HPO_4_ adjusted to pH 7.6). At 15 minutes, slides were sequentially dehydrated in 95% ethanol, 99% isopropanol, and CitriSolv (Decon Labs, #1601) to terminate the reaction, and were then mounted (Vectamount mounting media; Vector Laboratories), and cover-slipped. Slides were scanned (Motic EasyScan Pro 6 Digital Slide Scanner) and SDH activity was measured based on median pixel density across the entire muscle using a custom ImageJ (Fiji) macro.

### Mitochondrial respiration in frozen skeletal muscle

Respirometry was conducted on TA lysates, as previously described (39–41). Briefly, the TA was thawed in ice*-c*old PBS and then mechanically homogenized (10–20 strokes in a Teflon*-g*lass homogenizer) in MAS buffer (70 mM sucrose, 220 mM mannitol, 5 mM KH_2_PO_4_, 5 mM MgCl_2_, 1 mM EGTA, 2 mM HEPES pH 7.4). The homogenate was centrifuged (1,000 g, 10 min, 4°C) and the protein concentration of the supernatant determined by bicinchoninic acid assay (Thermo Fisher). Subsequently, tissue homogenates (6 μg) were loaded into a Seahorse XF96 microplate in 20 μL of MAS. The loaded plate was centrifuged (2,000 g, 5 min, 4°C, no brake) and 130 μL of MAS containing cytochrome c (10 μg/mL, final concentration), was added. Substrate injection was as follows – Port A: pyruvate + malate (5 mM each), NADH (1 mM), or 5 mM succinate (5 mM) + rotenone (2 μM); Port B: rotenone (2 μM) + antimycin A (4 μM); Port C: TMPD + ascorbic acid (0.5 mM + 1 mM); Port D: azide (50 mM). Oxygen consumption rate (OCR) normalized to protein was calculated using Seahorse Wave software (Agilent). Respiratory capacity through Complex I (CI), Complex II (CII), and Complex IV (CIV) was calculated by subtracting OCR values after injection of inhibitors from the substrate-induced maximal OCR (41).

### Microarray data

We interrogated published microarray data (26, 38) that studied circadian variability in gene expression in mouse skeletal muscle (GEO accession: GSE3746); for our analysis, raw intensity for genes of interest were extracted from the normalized data file (i.e., GSE3746_series_matrix file) in which significant circadian rhythmicity was assigned to all genes with a multiple measures corrected β (MMCβ) value ≤ 0.2. From this published data, the significant circadian genes list was prepared by converting the LocusLink to Emsembl geneID using DAVID (42) and ranked by the inverse of the MMCβ value. The enrichment of MitoPathways3.0 specified in the mitochondrial protein database Mouse.MitoCarta3.0 (43) gene sets was analyzed and visualized using the fgsea, clusterProfiler and ggplot2 packages in R.

### Sample blinding and statistics

After collected, all muscle samples were randomly numbered, with experimenters blinded to the group (i.e. ZT timepoint) assignment when conducting analysis. For the qPCR, immunoblotting, SDH activity, and uncoupled mitochondrial respiration data, assessment of circadian rhythmicity was determined using JTK_CYCLE (44). Statistical significance was set at a Adjusted P value of < 0.05. All data are expressed as mean ± SEM.

## Results

### Markers of mitochondrial biogenesis, homeostasis, and metabolism do not exhibit intrinsic circadian regulation in skeletal muscle

We measured relative mRNA expression of the core-clock gene *Bmal1*, the core-clock output gene *Dbp*, and a panel of genes involved in mitophagy (*Prkn, Bnip3, Pink1*), mitochondrial dynamics and mitochondrial DNA maintenance (*Fis1, Opa1*), and carbohydrate and lipid metabolism (*Pdha1, Ogdh, Ctp1b, Acadl*), in EDL, SOL, and GA collected at ZT 1, 7, 13, and 19. As expected, *Bmal1* and *Dbp* demonstrated circadian rhythmicity in all three muscles. Specifically, for *Bmal1*, the zenith was 6.0-, 20.2-, and 5.1-fold higher than the nadir in the EDL (Figure 1A), SOL (Figure 1B) and GA (Figure 1C), respectively. For *Dbp*, the zenith was 14.1-, 9.7- and 11.8-fold higher than the nadir in the EDL (Figure 1A), SOL (Figure 1B) and GA (Figure 1C), respectively. In contrast, none of the genes that we measured as markers of mitochondrial function were regulated by time-of-day (EDL: Figure 1A; SOL: Figure 1B; GA: Figure 1C). Similarly, none of these genes, (except for *Bmal1* and *Dbp*), were identified as circadian-regulated in the circadian transcriptome papers in skeletal muscle by McCarthy et al. (26) and Miller et al. (38).

**Figure 1.**
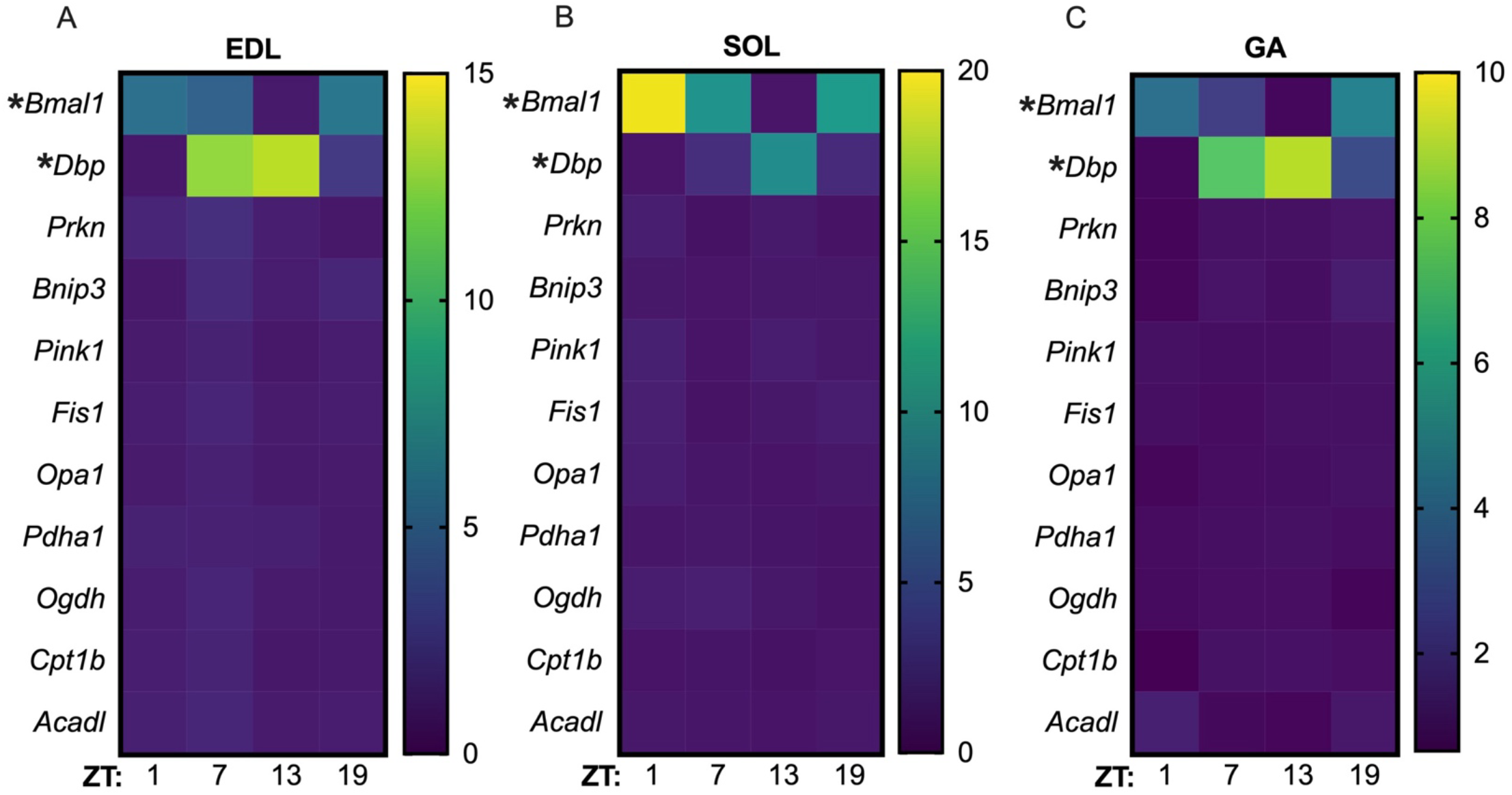
mRNA expression levels of genes involved in mitochondrial biogenesis, homeostasis, and function do not vary by time-of-day in skeletal muscle despite robust circadian rhythmicity of core-clock and clock-output genes. (**A**) Heat map of transcript abundance from qPCR results in EDL. (**B**) Heat map of transcript abundance from qPCR results in SOL. (**C**) Heat map of transcript abundance from qPCR results in GA. (**A-C**) Rows correspond to gene names and columns correspond to zeitgeber time (ZT). Statistics: (**A**-**C**) Values for individual cells in the heat map are the median values of *n* = 3-4 biological replicated per gene-timepoint per muscle. **\*** = Adjusted P value of < 0.05 determined by JTK_CYCLE (44).

### Transcript and protein abundance of electron transport chain (ETC) complexes do not display intrinsic circadian regulation in skeletal muscle

For insight into circadian rhythmicity of ETC complexes, we measured the relative mRNA expression and protein abundance of ETC subunits for *mt-Nd6* (complex I), *Sdhb* (complex II), *mt-Co1* (complex IV) and *Atp5f1a* (complex V), in the EDL, SOL, and GA (transcript only). At the gene level, in the EDL, there was no significant circadian rhythmicity for any of the genes. For SOL, *Sdhb* and *Atp5f1a* did not exhibit circadian rhythmicity, whilst *mt-Nd6* and *mt-Co1* did, with a 1.9-fold delta in *mt-Nd6* (ZT1 vs. ZT7) and 1.8-fold delta in *mt-Co1* (ZT19 vs. ZT7) comparing the zenith to the nadir (Figure 2B). For GA, there was no circadian variation in any of these genes (Figure 2C). Similarly, none of the individual genes were identified as circadian-regulated by McCarthy et al. (26) or Miller et al. (38), which also studied the GA. Moreover, pathway analysis of the identified circadian-regulated genes in McCarthy et al. (26) and Miller et al. (38) using MitoCarta did not identify any significantly enriched terms (Supplementary Figure 1). At the protein level, there was no time-of-day variation in MT-ND6, SDHB, MT-CO1 or ATP5F1A in the EDL (Figures 2D-H) or SOL (Figures 2I-M).

**Figure 2.**
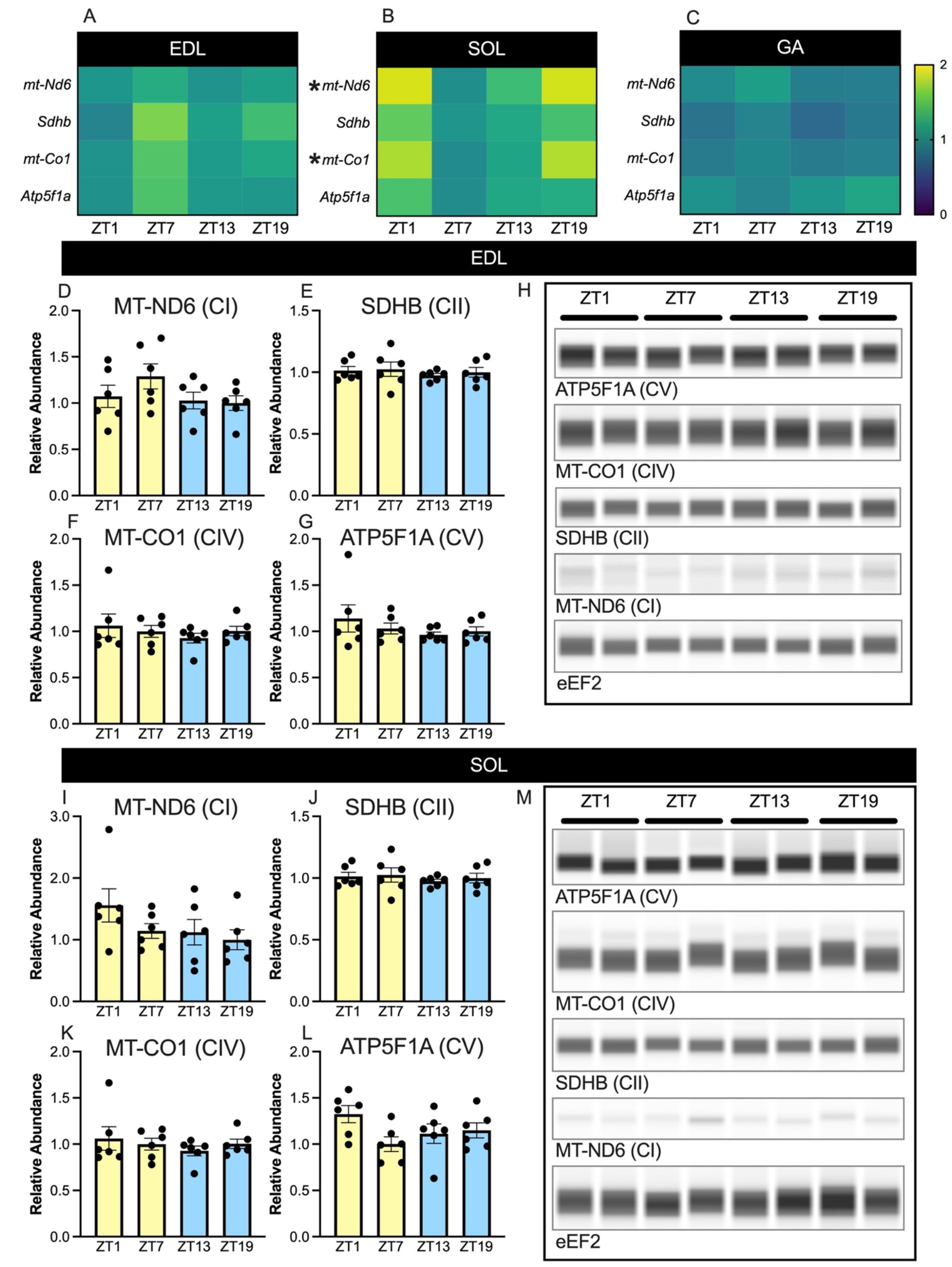
Abundance of electron transport chain proteins do not vary by time of day despite subtle circadian variation in mRNA expression in some skeletal muscles. (**A**) Heat map of qPCR results for mRNAs associated with different electron transport chain complexes (rows) at different times-of-day (zeitgeber time [ZT] 1, 7, 13, and 19) in extensor digitorum longus (EDL). (**B**) Heat map of qPCR results for mRNAs associated with different electron transport chain complexes (rows) at different times-of-day (ZT1, 7, 13, and 19) in soleus (SOL). (**C**) Heat map of qPCR results for mRNAs associated with different electron transport chain complexes (rows) at different times-of-day (ZT1, 7, 13, 19) in gastrocnemius (GA). (**D**) MT-ND6 (Complex I [CI]) protein abundance in EDL. (**E**) SDHB (Complex II [CII]) protein abundance in EDL. (**F**) MT-CO1 (Complex IV [CIV]) protein abundance in EDL. (**G**) ATP5F1A (Complex V [CV]) protein abundance in EDL. (**H**) Representative immunoblots for EDL protein abundance data. (**I**) MT-ND6 (Complex I [CI]) protein abundance in SOL. (**J**) SDHB (Complex II [CII]) protein abundance in SOL. (**K**) MT-CO1 (Complex IV [CIV])protein abundance in SOL. (**L**) ATP5F1A (Complex V [CV]) protein abundance in SOL. (**M**) Representative immunoblots for SOL data protein abundance data. (**D**-**G** and **I**-**L**) Blue bars indicate time-points during the dark phase and yellow bars indicate time-points during the light phase of the circadian cycle. Statistics: (**A**-**C**) Values for individual cells in the heat map are the median values of *n* = 3-4 biological replicates per gene-timepoint per muscle. (**D**-**G** and **I**-**L**) Data are expressed as mean ± SEM. **\*** = Adjusted P value of < 0.05 determined by JTK_CYCLE (44).

### Mitochondrial oxygen consumption rate and SDH activity do not exhibit intrinsic time-of-day regulation in skeletal muscle

For insight into circadian regulation of mitochondrial function itself, we measured maximal mitochondrial oxygen consumption in lysates from frozen TA. Thus, there was no effect of time-of-day on CI-, CII-or CIV-driven maximal respiratory capacity (Figure 3A). Similarly, there was no circadian variation in SDH (i.e. complex II) activity in the TA or PLN (Figure 3B-C and Supplementary Figure 2).

**Figure 3.**
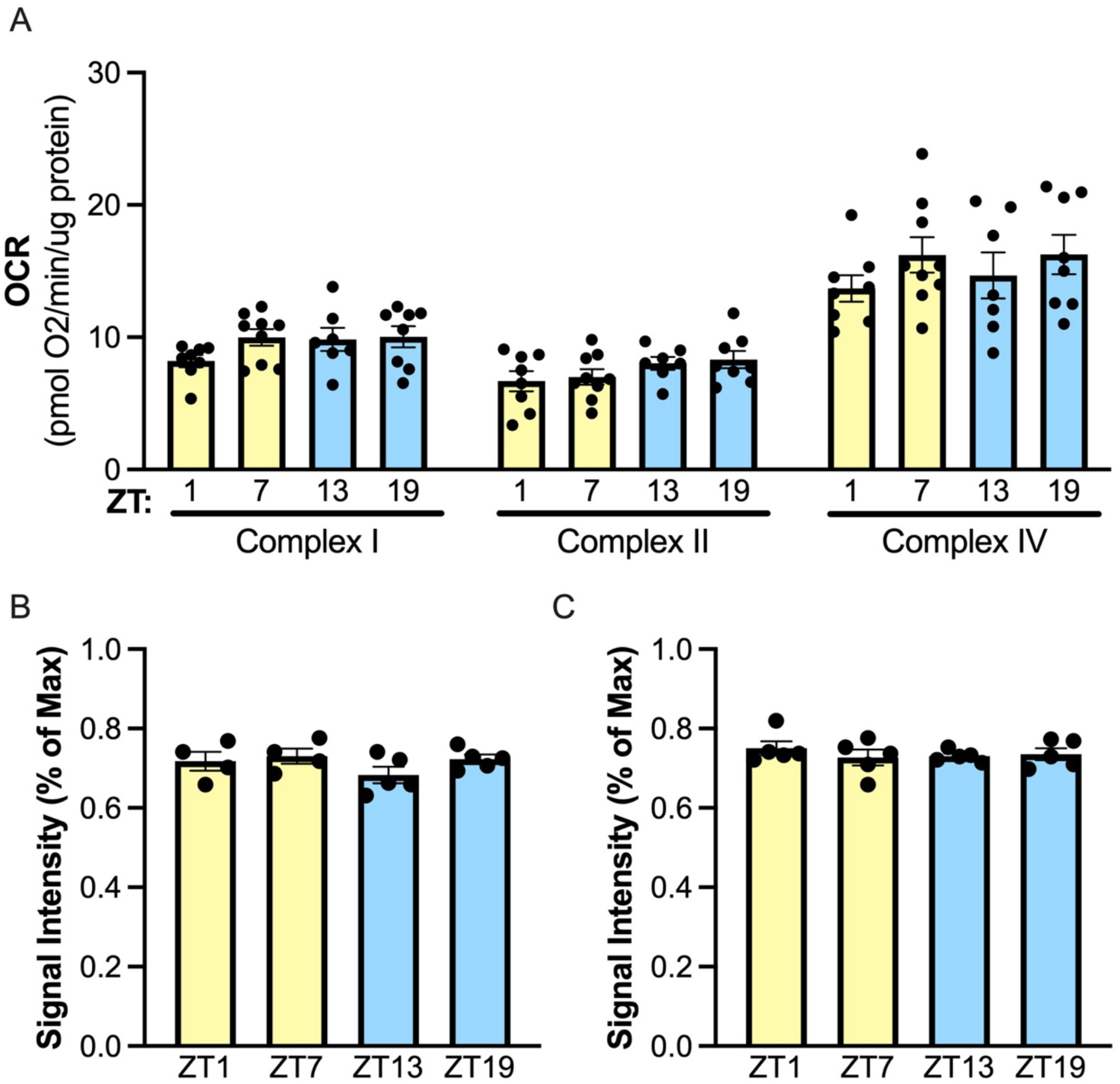
Maximum oxygen consumption and SDH activity are not regulated by time-of-day. (**A**) Oxygen consumption rate of tibialis anterior electron transport chain complexes at different times-of-day (zeitgeber time [ZT] 1, 7, 13, and 19). (**B**) Signal intensity of SDH activity expressed as a % of maximum intensity in tibialis anterior at 4 different times of day (zeitgeber time [ZT] 1, 7, 13, and 19). (**C**) Signal intensity of SDH activity expressed as a % of maximum intensity in plantaris at 4 different times of day (ZT1, 7, 13, and 19). Statistics: Data are expressed as mean ± SEM. No circadian rhythmicity was detected using JTK_CYCLE (44).

## Discussion

While markers of mitochondrial function are generally believed to be intrinsically regulated by time-of-day in skeletal muscle (13–17), whether markers of mitochondrial function across skeletal muscles are intrinsically regulated (i.e., is shared across different skeletal muscles and occurs independently of meal timing) remains to be fully defined. To this end, we measured mitochondrial function at the gene, protein, enzymatic, and functional level (i.e., O_2_ consumption) in muscle from post-absorptive female mice at four zeitgeber times (ZT), ZT 1, 7, 13, and 19. Overall, we found no evidence of circadian rhythmicity in markers of mitochondrial biogenesis, dynamics, or metabolism at the transcript (SOL, EDL and GA), protein abundance (SOL and EDL), or enzyme activity (PLN and TA) level. Moreover, maximal uncoupled mitochondrial respiration in the TA was not different by time-of-day. Taken together, these findings demonstrate that markers of mitochondrial function are not intrinsically regulated by time-of-day in female mouse skeletal muscle.

In 2007, two seminal papers (26, 38) established the circadian transcriptome of the GA in male C57BL6 mice. In these studies, 215 (38) or 267 (26) genes (depending on the multiple measures corrected β cut-off used) out of the 36,182 nonredundant mouse sequences analyzed demonstrated circadian rhythmicity. Interestingly, however, analyses done by the authors in the original papers (26, 38) and our own mitochondria-specific (using MitoCarta) enrichment analysis of their data failed to identify any significant mitochondrial terms. Similarly, we found no circadian rhythmicity in mitochondrial transcript expression in GA or EDL. While we did find small rhythmic changes in 2 ETC genes in SOL, since these were not shared between muscles, by definition they are not intrinsically regulated. This contrasts with a recent study in human skeletal muscle, which found time-of-day differences in gene expression of mitochondrial- and metabolic-related genes (31), including one of genes that we measured; *Cpt1b*. Species-related differences could explain the differences between these data in human muscle (31) and mouse muscle (26, 38). Alternatively, considering that a meal or acutely increasing systemic glucose and/or insulin availability profoundly impacts skeletal muscle gene expression (34–36), it is possible time-of-day effects in human muscle were due to the time elapsed since the last meal, rather than intrinsic circadian rhythmicity. For example, in one human study (31) a standardized meal was given before muscle biopsies during the light phase, but not during the dark-phase. Importantly, the lack of a time-of-day effect on mitochondrial and metabolic gene expression we observed was not related to our study design of controlling the last meal, as the core circadian genes, *Bmal1* and *Dbp*, were robustly regulated by time-of-day in the SOL, EDL, and GA. Similarly, feeding rats an ultradian diet (in which rats were fed 6 equally spaced meals per day) for 6 weeks minimally impacted core-clock gene rhythmicity, but did impact the rhythmicity of expression of metabolic genes (33). Thus, when considering our data and previously published data (26, 38) together, it is evident that common genes representative of mitochondrial biogenesis, dynamics, and metabolism are not intrinsically regulated by time of day in skeletal muscle.

The consensus view in the field of skeletal muscle circadian biology is that mitochondrial function oscillates by time-of-day (13–15, 17, 19). For example, immortalized muscle-like C2C12 myoblasts demonstrate an intrinsic 24-h rhythmicity of mitochondrial oxygen consumption, independent of serum exposure (i.e. feeding) (22); notably, however, this experiment was conducted in n = 1 at each timepoint, so it will be of interest to confirm this finding in future work. In germline KO mouse studies of important circadian proteins, maximal oxygen consumption is impaired in the permeabilized soleus from mice with global knockout of *Nr1d1*^-/-^ (Rev-Erbα) (25) and in the gastrocnemius and diaphragm from mice with global knockout of *Bmal1*^-/-^ mice (24). Furthermore, maximal fatty acid oxidation is higher in primary myotubes established from dKO^*Cy1/Cry2*^ mice compared to WT littermates (23). However, mitochondrial function throughout a 24-h cycle was not studied (i.e., measures were made at one timepoint). Also, the potential impact of core-clock proteins during muscle development and the subsequent effects on markers of mitochondrial function were not studied. To this point, *Bmal1*^*-/-*^ mouse embryonic stem cells display widespread dysregulation of ecto-, meso-, and endoderm differentiation (45, 46), occurring in conjunction with changes in mitochondrial function and substrate utilization (46). In contrast, *Bmal1*^*-/-*^ mice with post-developmental, whole-body KO of *Bmal1* in various tissues exhibit normal life span, fertility, body weight, and glycemia (47). Finally, mice with post-developmental *Bmal1* KO selectively in skeletal muscle do not exhibit any impairment in muscle function, skeletal muscle fiber type distribution, or skeletal muscle mitochondrial function (as measured by SDH activity) (48). In contrast, these measures are overtly changed in mice with germline KO of *Bmal1* (24, 49). Collectively, this demonstrates that the mitochondrial phenotype in germline *Bmal1*^-/-^ KO mice is most likely related to its role during fetal (muscle) development, rather than a fundamental role of the circadian clock in skeletal muscle contractile or mitochondrial function.

In humans, mitochondrial state 3 respiration (20, 31, 50) may display time-of-day regulation under energy demanding conditions in muscle. Interestingly, this time-of-day variation is independent of mtDNA copy number, ETC protein abundance, citrate synthase activity, or mitochondrial voltage-dependent anion channel protein abundance, which collectively do not display circadian rhythmicity in human (20, 50) and rat (29) muscle. Considering the impact of acute changes in systemic glucose and insulin availability (51) or timing of feeding (29) on skeletal muscle mitochondrial function, it will be of interest in future studies to determine if the effects on mitochondrial respiration (20, 31, 50) are due to meal timing or are reflective of intrinsic circadian regulation. It is important to note that an effect of time-of-day on mitochondrial function in human muscle is not a universal finding (32, 50). A proposed reason for the lack of effect in these human studies is that the subjects were aged, overweight, and/or obese as compared to young subjects studied by van Moorsel et al. (20). Nevertheless, there was still no circadian rhythmicity in mitochondrial respiration after robustly improving mitochondrial function and glycemic control through 12 weeks of exercise training (32). Similarly, we found no effect of time-of-day on maximal oxygen consumption in the TA and no time-of-day regulation in SDH activity in the TA or PLN. In addition to controlling for the timing of the last meal, an advantage of our analytical approach is that we assessed mitochondrial oxygen consumption in muscles from each timepoint in the same analytical run, thus avoiding potential day-to-day variability issues (especially as it relates to sample preparation) that can impact measurement of mitochondrial oxygen consumption in skeletal muscle (52). In sum, these data suggest that markers of skeletal muscle mitochondrial function are not intrinsically regulated by time-of-day.

MHC composition can widely vary in a mixed skeletal muscle, such as the vastus lateralis, in humans (53–55). For example, a study that examined 418 human subjects found that the proportion of type I fibers ranged from 15% to 85% within the muscle and was accompanied by a large variation in markers of aerobic oxidative markers (53). In line with this, within-subject variation of type II and I fiber type distribution in multiple biopsies of the vastus lateralis was found to be ~18% (56), which is similar to the ~20% difference in mitochondrial oxidative capacity that is found by time-of-day in healthy human muscle (20). Thus, in future studies it will be important to compliment mitochondrial functional measures with measures of fiber type to ensure differences are not driven by the natural fiber type variability seen in vastus lateralis biopsies. Further to this point, we found quite different responses in circadian gene rhythmicity in muscles of widely different MHC composition. For example, in the EDL (which is exclusively fast twitch fibers in the mouse, comprising ~70-80% type IIb (57, 58) *Bmal1* changed by 6.0-fold, whilst in the SOL (which is ~30-40% slow twitch [type I] fibers and ~50% type IIa (57, 58)), *Bmal1* was changed by 20.2-fold. To our knowledge, no studies to date have investigated the impact of MHC composition on rhythmicity of circadian genes. Such an effect should be an important consideration in future studies, especially human studies given the variability within the vastus lateralis.

In conclusion, in line with our previous work showing that skeletal muscle endurance capacity is not impacted by time-of-day (37), our results demonstrate that markers of mitochondrial function, be it at the transcript, protein, enzymatic activity, or functional level, do not exhibit intrinsic circadian regulation in female mouse skeletal muscle. Interestingly, we found that the amplitude circadian rhythmicity of core-clock genes is differentially regulated across skeletal muscles, which may explain some of the inter-subject variability in human studies using the vastus lateralis. Moreover, considerations for future circadian studies in skeletal muscle include performing analyses in multiple muscles and controlling for food intake. Since circadian studies to date have almost exclusively studied males, this work in female mice is an important addition to the field of circadian biology in skeletal muscle.

## Acknowledgements

This work was supported, in part, by National Institutes of Health (NIH) grants R21 AR072882 and R21 AG067495 to S. Schenk, whilst L.S. Fitzgerald, N. Lau, and M. Melamed were supported, in part, by a Summer Research Fellowship from the UC San Diego School of Medicine. L.S. Fitzgerald was also supported by the NIH-funded UC San Diego Medical Scientist Training Program (T32 GM007198). C.R. Spurrier was supported by the Dr. Maryam Ahmadian Memorial Summer Fellowship Program at US San Diego. The authors also acknowledge support from the Wu-Tsai Human Performance Alliance and the Joe and Clara Tsai Foundation. L.A. Burnett was supported by the Reproductive Scientist Development Program (K12HD000849). A.L. Hevener is supported in part by funding from the UCLA Department of Medicine and NIH grant U54DK120342. We are also grateful for the support of the NIH-funded (P30 DK063491) UCSD-UCLA Diabetes Research Center, especially the Metabolic and Molecular Physiology Core (MMPC) and the Mitochondrial Biology sub-core, which is directed by Drs. Orian Shirihai and Linsey Styles.

## Author Contributions

G.A.M. and S.S. conceived and designed research; L.S.F, C.R.S., N.L., M.M., and C.G. performed experiments; L.S.F., C.R.S., G.A.M., C.G., A.L.H., J.A.S., and S.S. analyzed data; L.S.F, C.R.S., N.L., M.M., L.A.B., G.A.M., C.G., A.L.H., J.A.S., and S.S. interpreted results of experiments; L.S.F., C.G., G.A.M., J.A.S., and S.S. prepared figures; L.S.F. drafted manuscript; L.S.F. and S.S. edited and revised manuscript; L.S.F, C.R.S., N.L., M.M., L.A.B., G.A.M., C.G., A.L.H., J.A.S., and S.S. approved final version of manuscript.

