## Supplementary Table 1 and Supplementary Figures 1 and 2 for "Markers of mitochondrial function and oxidative metabolism in female skeletal muscle do not display intrinsic circadian regulation"

**Supplementary Table 1.** Primers for semi-quantitative RT-PCR analysis.

| **Gene Name** | **FWD** | **REV** |
|---|---|---|
| *Bmal1* | CACTGTCCCAGGCATTCCA | TTCCTCCGCGATCATTCG |
| *Dbp* | CCTGAGGAACAGAAGGATGA | ATCTGGTTCTCCTTGAGTCTT |
| *Prkn* | TCAAGAAGACCACCAAGCCT | AACCAGTGATCTCCCATGCA |
| *Bnip3* | GGGCTCCTGGGTAGAACTG | GGGCTCCTGGGTAGAACTG |
| *Pink1* | CGAGCATCTTCTAGCCCTGA | CGAGCATCTTCTAGCCCTGA |
| *Fis1* | AGAGGAACAGCGGGACTATG | AGAGGAACAGCGGGACTATG |
| *Opa1* | ACAAGATCACCTACCACGGG | ACAAGATCACCTACCACGGG |
| *Pdha1* | ATGGGGACGTCTGTTGAGAG | ATGGGGACGTCTGTTGAGAG |
| *Ogdh* | GGTGGAAGCACAACCTAACG | GGTGGAAGCACAACCTAACG |
| *Cpt1b* | ATCTGGGCTATCTGTGTCCG | ATCTGGGCTATCTGTGTCCG |
| *Acadl* | ATCTGGGCTATCTGTGTCCG | TAAGTCACTTCCAGCCCCAG |
| *mt-Nd6* | GGGATGTTGGTTGTGTTTGGA | GGGATGTTGGTTGTGTTTGGA |
| *Sdhb* | TGTACGAGTGCATCCTGTGT | CGGTAGACAGAGAAGGGGTC |
| *mt-Co1* | CGGTAGACAGAGAAGGGGTC | GTCAGTTTCCAAAGCCTCCA |
| *Atp5f1a* | CTGCCACTCAACAGCTCTTG | GTGCTGGCTGATAACGTGAG |
| *Rplp0* | GTTTGACAACGGCAGCATTT | GCGCTTGTACCCATTGATGA |

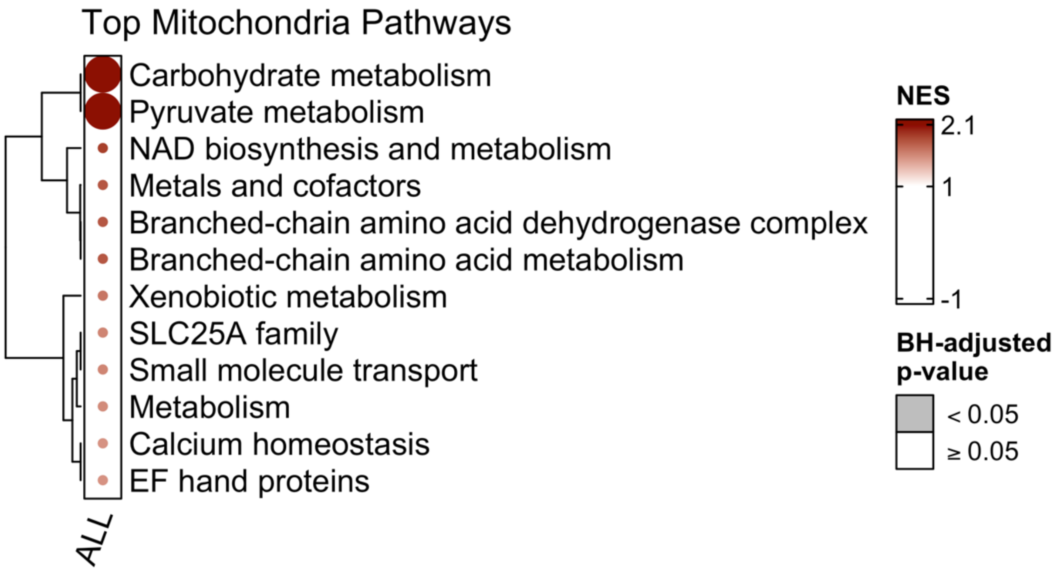


**Supplementary Figure 1. Mitocarta pathway analysis of previously published microarray data does not demonstrate time-of-day enrichment of mitochondrial pathways**. We accessed (GEO accession: GSE3746) previously published microarray data from McCarthy et al. (26) and Miller et al. (38) that identified the circadian transcriptome of mouse skeletal muscle. Genes from this analysis were ranked using the inverse of multiple measures correct β (MMCβ) which indicates goodness of fit to a cosine wave form over a ≈24-h period. Ranked genes were then analyzed using the mitochondrial protein database Mouse.MitoCarta3.0 to identify enriched mitochondrial pathways. No mitochondrial pathways were significantly enriched in this analysis ($p \geq 0.05$).

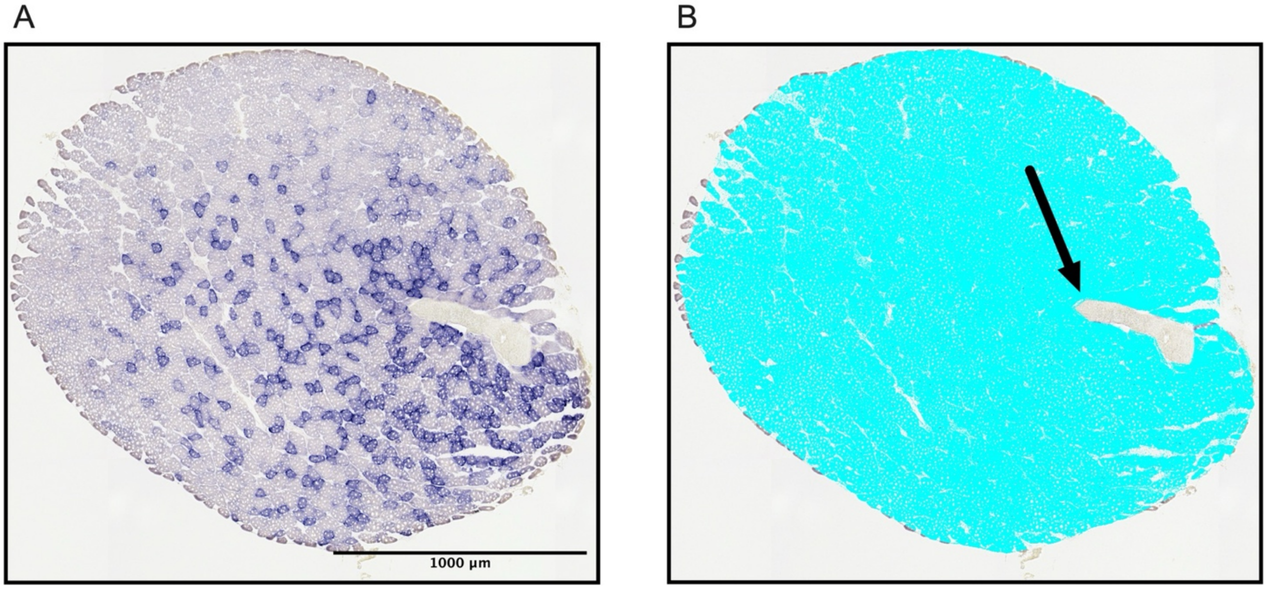


**Supplementary Figure 2. Representative image of skeletal muscle stained for assessment of succinate dehydrogenase (SDH) activity.** (**A**) Example raw image of a 10 µm section of plantaris used for assessment of SDH activity. (**B**) Example of how area on the slide was limited to exclude empty glass, tendon, and interfibrillar space; quantified area is highlighted in blue. Arrow points to tendon.
